# Immune reprogramming of 3D tumor models via optoporation-mediated targeted gene delivery to macrophages

**DOI:** 10.64898/2026.03.16.712123

**Authors:** Ines Poljak, Izzah Nabilah Hussein, Chenlei Gu, Giulio Giustarini, Teng Xiang, Yusuke Toyama, Ciro Chiappini, Giulia Adriani

**Affiliations:** Centre for Craniofacial and Regenerative Biology, King’s College London, London, United Kingdom; Singapore Immunology Network (SIgN), Agency for Science, Technology and Research (A*STAR), Republic of Singapore; A*STAR Skin Research Labs (A*STAR SRL), Agency for Science, Technology and Research (A*STAR), Republic of Singapore; London Centre for Nanotechnology, King’s College London, London, United Kingdom; Mechanobiology Institute, National University of Singapore (NUS), Republic of Singapore; Department of Biomedical Engineering, National University of Singapore (NUS), Republic of Singapore; Lee Kong Chian School of Medicine, Nanyang Technological University, Republic of Singapore

## Abstract

The dynamics of the tumor microenvironment (TME) are a key determinant of cancer progression and therapeutic resistance through complex interactions between tumor, stromal and immune cell populations. Among these, tumor-associated macrophages (TAMs) play a central role in promoting tumor growth and immune suppression. However, the specific contributions of TAMs remain poorly understood due to the lack of tools enabling selective genetic manipulation in three-dimensional (3D) tumor models. Here, we present a gold nanoparticle-assisted optoporation approach that enables spatially selective plasmid-based gene delivery to TAMs within intact heterocellular 3D pancreatic ductal adenocarcinoma (PDAC) spheroids, thereby modulating the TME. In two-dimensional (2D) TAM cultures, conventional transfection of *IRF5*- and *IKBKB*-encoding plasmids validated their capacity to induce TAM repolarization, as evidenced by activation of interferon signaling. Extending this approach to 3D PDAC spheroids, nanoparticle-assisted optoporation achieved selective transfection of TAMs with *IRF5-* and *IKBKB-*encoding plasmids by transiently generating nanoscale membrane pores in illuminated cells. TAMs transfection elicited a robust interferon response, marked by transcriptional upregulation of *IFNA, IFNB1*, and *CXCL10*, and increased protein levels of IFNB1, IFNL1, and CXCL13, together with downregulation of pro-tumorigenic markers CEACAM5, IL19, and IL32. These coordinated changes indicate a shift towards an anti-tumorigenic TME. By enabling minimally invasive, TAM-specific gene delivery in complex multicellular 3D spheroids, this strategy allows precise modulation of the TME and opens new avenues for modeling its dynamics in cancer progression and therapeutic response.

## Main

The tumor microenvironment (TME) is a dynamic and heterogeneous network, composed of cancer, stromal and immune cells embedded in a complex extracellular matrix and regulated by diverse mechanical and biochemical cues. These reciprocal interactions influence tumor initiation, progression, metastasis and therapeutic response^1–3^. Therefore, replicating the functional diversity and complex molecular communication among cells *in vitro* is essential for better understanding of the tumor biology and therapy development. An important factor driving TME dynamics is the polarization of tumor-associated macrophages (TAMs) toward pro-tumorigenic phenotypes, which is largely driven by tumor cells and microenvironmental cues. Polarized TAMs, in return, promote tumor immune evasion and elicits paracrine signaling, further reshaping the microenvironment^4^.

This mutual interplay is particularly evident within pancreatic ductal adenocarcinoma (PDAC) where TAMs play a central role in influencing tumor behavior, providing a biologically relevant target for modulation^5–7^. PDAC, a third leading cause of mortality in cancer patients^8^, is characterized by suppressed type I interferon (IFN) signaling^8^. Type I IFNs exert potent anti-tumor effect through both direct mechanisms, including tumor cell apoptosis, and indirect mechanisms, involving recruitment of natural killer (NK) and B cells^9^. Consistently, enhanced IFN signaling has been linked to improved survival in PDAC mouse models^10^. Interferon regulatory factor (IRF5) and IkappaB kinase (IKKβ), two master regulators of pro-inflammatory and IFN signaling^11–14^, have been shown to induce anti-tumor macrophage activation in mouse models of ovarian cancer and melanoma^15^. However, whether localized IRF5–IKKβ–driven macrophage reprogramming can overcome pro-tumorigenic TAM polarization and propagate a TME remodeling in PDAC has not been addressed. To address this gap, we cultured humanized PDAC spheroids comprising pancreatic cancer cells (PANC-1), pancreatic stellate cells (PSCs), endothelial cells (ECs), and blood-derived monocytes that organize spontaneously into a tissuelike structure modeling key features of patients TME including TAM generation^16^. This physiologically relevant model provides an opportunity to rapidly probe immune-tumor crosstalk under near-native conditions, enabling direct manipulation of defined cell populations, namely TAMs, within a 3D microenvironment and assessment of whether TAM reprogramming toward an anti-tumor state can propagate functional changes across the TME^17^. Achieving this, however, requires tools that enable cell-type-specific genetic perturbation tools in complex 3D tumor models.

Gene delivery by transfection provides a powerful method for perturbing molecular networks to study cellular function. Conventional approaches such as lipofection or electroporation, although applicable to complex 3D TME models^18,19^, lack the spatial precision required for singlecell manipulation, affecting all cells and limiting the ability to identify the specific cellular source of signals during therapy response or disease progression. Selective manipulation of individual cells *in situ* is instead essential for modeling dynamic changes within the TME. Nanoparticle-mediated optoporation addresses this limitation by combining structured light and nanoparticle-driven energy conversion to transiently open cell membranes for targeted gene delivery^20–25^. Optoporation efficiently transfects selected cells in 2D cultures with minimal toxicity^23,26,27^, and emerging approaches now enable transfection of 3D spheroids and organoids^21^. Thus, optoporation represents a promising strategy for selective TME modulation within 3D models through single-cell manipulation.

Here we establish an optoporation approach for the selective reprogramming of TAMs within PDAC spheroids by targeted *IRF5-* and *IKBKB-*encoding plasmid transfection. We demonstrated that nanoparticle-assisted optoporation enables TAM-specific gene delivery and functional reprogramming in intact 3D heterocellular spheroids. TAMs were pre-labeled with a fluorescent marker prior to spheroid formation to increase targeting specificity, and co-cultured with gold nanoparticles (AuNPs) to reduce the required laser power and overall toxicity. Following plasmid addition, the labelled TAMs were selectively illuminated, resulting in transient membrane poration exclusively in the targeted cells, permitting extracellular plasmid entry and transfection. Remarkably, targeted perturbation of TAMs led to an increase in interferons α (*IFNA)*, interferon β1 (*IFNB1)*, and interferon gamma-induced protein 10 *(CXCL10)* gene expression and IFNβ1, interferon λ1, (IFNλ1) and B lymphocyte chemoattractant (CXCL13) protein levels across the spheroid, demonstrating that localized immune activation can propagate through the entire TME.

This platform enables spatially precise, minimally invasive targeted transfection within intact 3D heterocellular spheroids for real-time *in situ* genetic manipulation of defined cell populations, overcoming the long-standing challenge of cell-type-selective transfection in multicellular environments. By reprogramming TAMs within PDAC spheroids, our approach allows investigating how TAM perturbations trigger a system-wide immune activation, establishing a broadly applicable framework for dissecting cell-specific mechanisms in complex tissues and their impact on TME reprogramming.

## Results

### Optoporation of human PBMC-derived TAMs

We first assessed the technical feasibility and biological suitability of TAMs for nanoparticle-assisted optoporation. We generated TAMs from human peripheral blood mononuclear cell (PBMC)-derived monocytes cultured in PANC-1 conditioned media^28^, and assessed their viability following exposure to AuNPs of 100 nm or 200 nm diameter, across a concentration range (Fig. 1a). Although modest, 100 nm AuNP caused a concentration-dependent decrease in viability beginning at 6 μg/mL, reaching statistical significance at 16 μg/mL (Fig. 1a, top panel). In contrast, 200 nm AuNPs did not affect TAMs viability across all tested concentrations (Fig. 1a, bottom panel). The difference between 100 and 200 nm AuNPs was observed at concentration of 6 μg/mL and reached significance at 12 μg/mL (Supplementary Fig. 1a). Accordingly, we selected 200 nm AuNPs at 6 μg/mL for all subsequent experiments.

**Figure 1.**
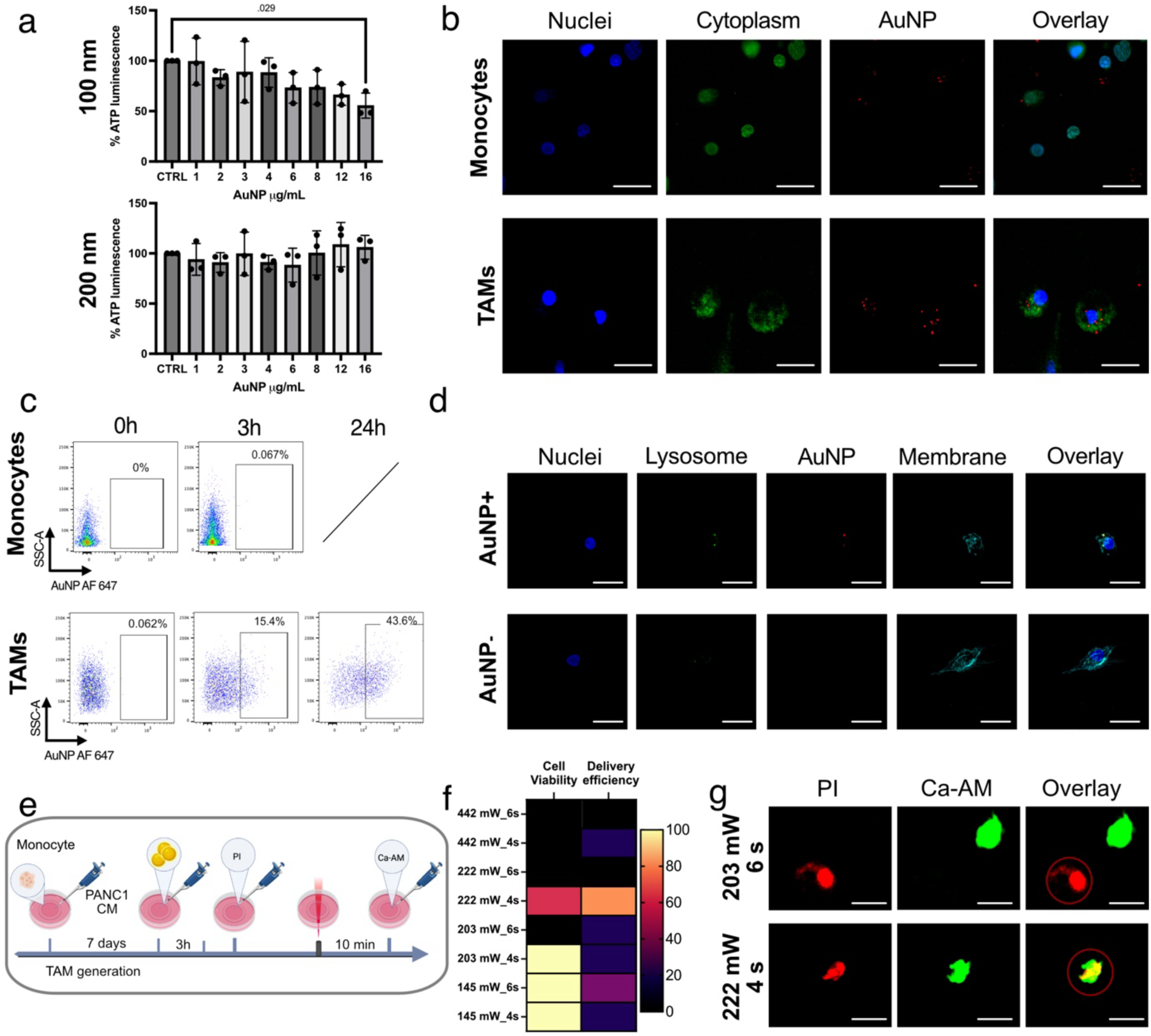
Optimization of gold nanoparticle (AuNP)-assisted optoporation parameters for gene delivery into human tumor associated macrophages (TAMs) in 2D cultures. **a,** Quantification of ATP-based cell viability in TAMs following 3 h incubation with or without (CTRL) 100 nm (top panel) and 200 nm (bottom panel) AuNPs at increasing concentration. Data represent mean ± SD from three biological replicates (n=3). **b,** Confocal images of human PBMC-derived monocytes (top panel) and TAMs (bottom panel) following 3 h of incubation with fluorescently labelled AuNPs (in red). Cell nuclei are in blue and cell cytoplasm is in green. **c,** Flow cytometric quantification of AuNPs uptake in monocytes and TAMs at 0 h, 3 h and 24 h post-incubation. **d,** Subcellular localization of AuNPs in TAMs with lysosomal marker staining. **e,** Schematic representation of the experimental design for testing the optoporation efficiency in 2D. **f,** Quantification of cell viability and delivery efficiency from confocal images with TAMs optoporated at different energy/pulse and speed. **g,** Representative confocal images of membrane permeabilization (propidium iodide in red) and viable cells (Calcein-AM in green) in TAMs at 203 W, 6 s (top panel) and 222 W, 4 s (bottom panel). Targeted cells are circled in red. Statistical analysis was done by oneway ANOVA with Dunnett’s test for multiple comparisons; p ≤ 0.05 was considered significant. Scale bars: 25 μm. Scheme in a and e was created with BioRender.

We next evaluated AuNP uptake efficiency by TAMs in comparison with undifferentiated monocytes. Confocal microscopy revealed increased uptake of AuNPs by TAMs compared to the undifferentiated monocytes (Fig. 1b). Flow cytometry confirmed the substantially greater uptake by TAMs with an uptake of 15.4% at 3 h, increasing to 43.8% at 24 h, while a minimal uptake was observed in the monocytes with 0.06% at 3 h and uptake at 24 h not assessed due to limited monocyte survival in culture (Fig. 1c). AuNPs were clearly observed within TAMs with high-resolution AiryScan imaging (Supplementary Figure 1b), while subcellular localization analysis demonstrated that AuNPs were predominantly localized within lysosomes of TAMs, as evidenced by the strong spatial overlap between the AuNP signal and lysosomal staining (Fig. 1d).

To optimize optoporation efficiency we evaluated propidium iodide (PI) delivery as an indicator of membrane permeabilization followed by Calcein-AM (Ca-AM) viability staining (Fig. 1e). We observed that higher laser power enhanced membrane poration efficiency but reduced cell survival, while with lower laser power cells were more viable but less porated. Optimal optoporation condition was achieved with 222 W power delivered in 4 seconds (Fig. 1f), yielding 80% optoporation efficiency with 60% post-optoporation cell viability (Fig. 1g). These 2D optimization data show that nanoparticle-assisted optoporation is a biologically compatible strategy for gene delivery into human TAMs and, we therefore applied these parameters in all subsequent experiments in more complex 3D spheroid models (Supplementary Table 1).

### Formation of AuNP-loaded PDAC spheroids

To establish a physiologically relevant 3D model of the PDAC TME, we generated heterocellular spheroids by co-culturing PANC-1 cancer cells, PSCs, ECs and PBMC-derived monocytes, herein referred to as naive spheroids, in which monocytes were incorporated during assembly and spontaneously polarized into TAMs through tumor-stromal interactions^16^ (Fig 2a). Here, we leverage scRNA-seq data from monocyte-seeded PDAC spheroids to address whether these models are suitable for studying TAM-driven TME remodeling. A broader, descriptive multimodal analysis of cellular complexity in 3D PDAC spheroids has been reported previously^29^. The scRNA analysis resolved distinct cancer cell, fibroblast, endothelial cell, and macrophage clusters, confirming the heterocellular composition of the 3D model (Fig. 2b). A supervised sub-clustering revealed five transcriptionally distinct TAM subtypes with characteristic gene signatures and associated biological processes (BP): inflammatory-associated (IA) TAMs expressing *IL1B*, *TNFAIP3* with enriched pathways in LPS-immune response and adaptive immunity; lipid-associated (LA) TAMs expressing *APOE*, *APOC1* with enrichment in lipoprotein and cholesterol metabolism; chemotaxis-associated (CA) TAMs expressing *CCL2*, *CXCL3*, *CCL4* with enrichment in chemotaxis and cell migration; metabolically active (MA) TAMs expressing *TXNIP* with enrichment in lipoprotein and cholesterol metabolism, and MAPK inhibition; and fibrosis-associated (FA) TAMs expressing *PDK4*, *F13A1*, *S100A9* with enrichment in chemotaxis and phagocytosis (Fig. 2c,d). This layered TAM diversity demonstrates that the 3D PDAC spheroid system recapitulates multiple functionally specialized macrophage states, providing a mechanistically relevant framework for testing TAM-targeted genetic reprogramming interventions. Analysis of *IRF5* expression revealed low expression levels across all TAM subsets, with the highest expression in the FA TAM population (Fig. 2e, Supplementary Fig. 2a). This low baseline *IRF5* expression is well suited for our IRF5-mediated TME remodeling strategy, as it allows broad targeting of TAMs without subset-specific restriction. Notably, *IRF5* expression did not correlate with *IL1B* expression (Fig. 2e), a cytokine associated with poor prognosis in PDAC patients, suggesting that the IRF5-driven reprogramming engages distinct, IL1B-independent signaling pathways that may contribute to tumor suppression.

**Figure 2.**
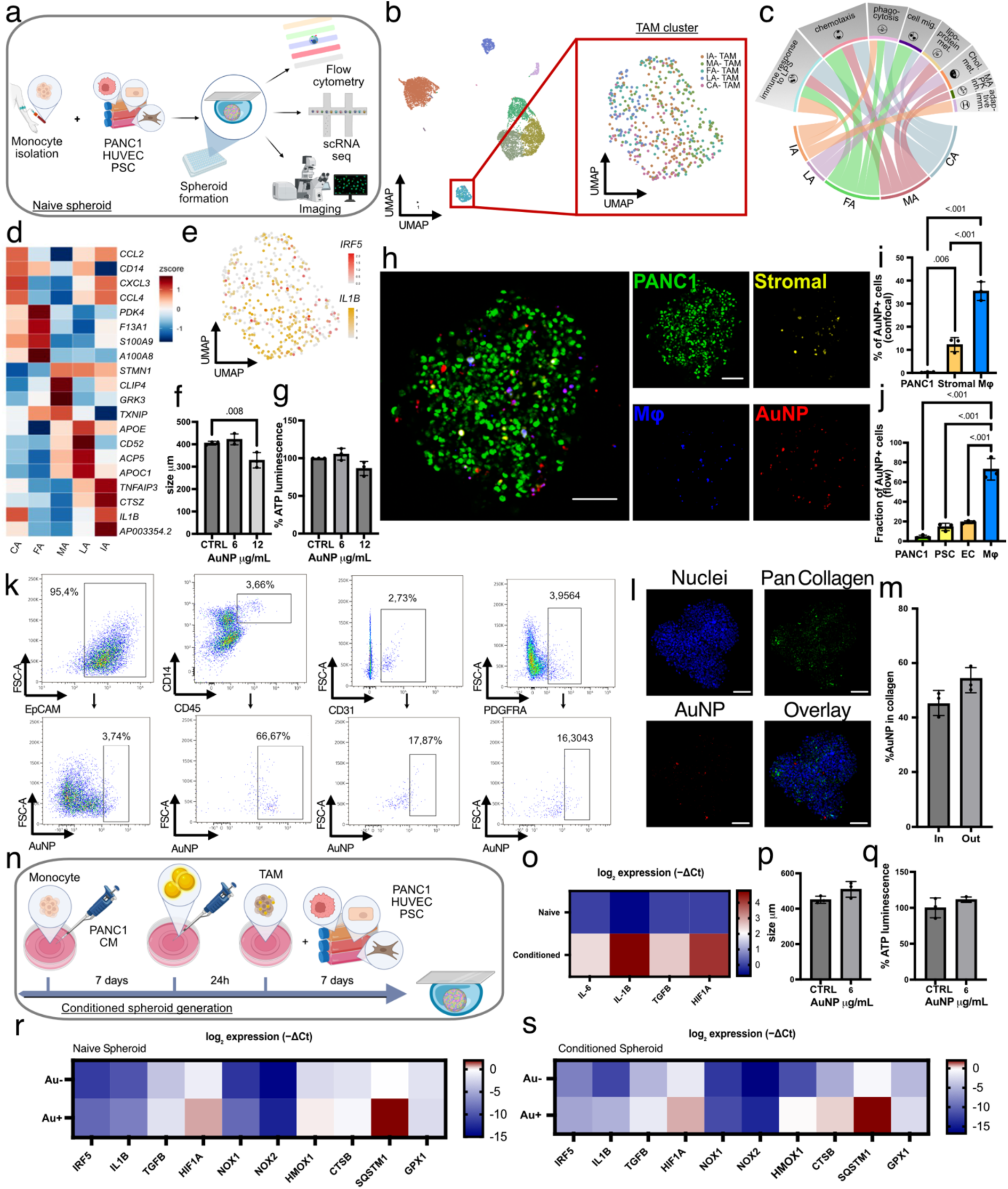
Characterization of TAM heterogeneity and optimization of AuNP incorporation in heterocellular PDAC spheroids for subsequent TAM-targeted optoporation. **a,** Schematic of the naive spheroid generation, where monocytes differentiate into TAMs *in situ*. **b,** UMAP of clusters in naive spheroids with magnified macrophage subclustering. **c**, Chord diagram of selected enriched biological process (BP) pathways in five TAM subtypes: inflammatory-associated (IA), lipid-associated (LA), chemotaxis-associated (CA), metabolically active (MA), and fibrosis-associated (FA) TAMs. **d,** Heatmap of the top 5 differentially expressed genes (DEGs) identified in each TAM subtype. **e,** Feature plot of *IRF5* and *IL1B* co-expression within the macrophage cluster. **f,** Size measurement of monocyte-seeded spheroids w/wo AuNP (6, 12 µg/mL; CTRL). **g,** Quantification of ATP-based viability in naive spheroids w/wo AuNP (6, 12 µg/mL; CTRL). **h,** Maximum intensity projection of the confocal image of naive spheroid formed with fluorescently labelled AuNPs, labelled for macrophages, stromal and cancer cells. **i,** Quantification of percentage of AuNP^+^ cells within each population by imaging analysis, PSCs and ECs are grouped as stromal cells and represented by the same color, AuNP at 6 µg/mL. **j,** Flow cytometric quantification of AuNP retention by each population at 6 µg/mL AuNP concentration. Cells were first gated on the individual cell population, followed by gating on AuNP⁺ cells. The y-axis shows the fraction of AuNP⁺ cells within each gated population. **k**, Representative flow cytometry gating strategy for data quantification presented in j. **l,** Maximum intensity projection of a representative confocal image of naive spheroid formed with 6 ug/mL fluorescently labelled AuNP, immunostained with anti-Pan-Collagen antibody. **m,** Quantification of AuNPs localized within (In) or outside (Out) collagen-rich extracellular matrix regions by imaging analysis. **n,** Schematic of conditioned spheroid generation, where pre-differentiated, AuNP-loaded TAMs are incorporated during spheroid assembly. **o,** qPCR quantification (log₂ expression, −ΔCt) of *IL6, IL1B, TGFB, HIF1A* in conditioned compared to naive spheroids. **p,** Size measurement of conditioned spheroids w/wo AuNP (6 µg/mL; CTRL). **q,** Quantification of ATP-based viability in conditioned spheroids formed w/wo AuNP (6 µg/mL; CTRL). **r,** qPCR quantification (log₂ expression, −ΔCt) of selected interferon-, inflammatory-, and stress-response genes in naive spheroids w/wo AuNP (6 µg/mL; CTRL). **s,** qPCR quantification (log₂ expression, −ΔCt) of gene expression of the same gene panel in conditioned spheroid w/wo AuNP (6 µg/mL; CTRL). Data represent mean ± SD from three biological replicates (n=3). Statistical analysis was done by one-way ANOVA with Dunnett’s test for multiple comparisons (**f, g, i, j**) or two-tailed unpaired Student’s t-test (**m, p, q**); p ≤ 0.05 was considered significant. Scale bars: 100 μm. Schemes in **a** and **n** were created with BioRender.

Next, we formed naive spheroids in the presence of AuNPs, further referred to as naive AuNP^+^ spheroids. We assessed spheroid size and viability following formation in the presence of 6 or 12 µg/mL AuNPs. While 6 µg/mL did not affect spheroid size or viability, spheroids formed with 12 µg/mL AuNPs exhibited reduced spheroid size. Therefore, 6 µg/mL AuNPs were selected for all subsequent experiments (Fig. 2f,g). Confocal imaging demonstrated that AuNPs were incorporated within the spheroid and predominantly localized to TAMs (Fig. 2h,i). While a distinct stromal core organization, as described in other four-cell spheroid models^16^, was not observed in this system, the images illustrate the heterocellular composition of our spheroids and the cell spatial distribution (Fig. 2h). Flow cytometry analysis confirmed that TAMs were the primary AuNP-retaining population (63.67% on average), followed by ECs (17.27%), PSCs (10.01%), and PANC-1 cells (7.17%) (Fig. 2j,k, gating strategy in Supplementary Fig. 2b). UMAP embedding of flow cytometry data showed that AuNP-associated signal co-localized with CD14⁺ cells, further suggesting preferential nanoparticle uptake by the macrophage population (Supplementary Fig. 2c). However, this approach of incorporating AuNP during spheroid formation yielded significant AuNP sequestration within the extracellular matrix (ECM). Staining with anti–pan-collagen antibody confirmed that on average 45.39% of AuNP were co-localized with collagen-rich regions (Fig. 2l,m), limiting bioavailability for optoporation.

Because AuNPs added directly to spheroids become trapped within the extracellular matrix (ECM), we pre-loaded nanoparticles into monocytes that had been pre-differentiated using PANC-1–conditioned media (TAMs) prior to spheroid assembly, aiming to improve their bio-availability for targeted optoporation. The resulting spheroids generated with pre-differentiated TAMs were termed conditioned spheroids, in contrast to naive spheroids, which were generated from undifferentiated PBMC-derived monocytes (Fig. 2n). Preferential AuNP retention in TAMs, but not monocytes, was previously confirmed (Fig. 1a,b,c). To validate the biological relevance of TAM-seeded spheroids as a PDAC TME model, we compared gene expression profiles between conditioned and naive spheroids without AuNPs. Conditioned spheroids displayed higher levels in gene expression of *IL6*, *IL1B*, *TGFB* and *HIF1A* (Fig. 2o), transcriptional markers consistent with features observed in PDAC patient samples, supporting their use as a relevant model^30–33^. The elevated cytokine levels may also reflect differences in macrophage number and/or activation state within the conditioned spheroids. We then formed conditioned spheroids with or without 6 μg/mL AuNPs present in TAMs and observed no differences in spheroid size (Fig. 2p) or viability (Fig. 2q). However, the presence of AuNPs increased major inflammatory and hypoxia-associated genes such as *TGFB*, *HIF1A*, *HMOX1*, *CTSB* and *SQSTM1* in both naive and conditioned spheroids, with a similar pattern but slightly higher levels in conditioned spheroids suggesting they could be more responsive to AuNP-mediated stress (Fig. 2r,s).

Together, these data show that incorporating AuNP loaded TAMs into heterocellular PDAC spheroids preserves spheroid integrity and viability while enriching nanoparticles within macrophages, providing a configuration that can be tested for a more selective TAM targeted optoporation in 3D.

### TAM-targeted optoporation reshapes the TME

To first determine whether transfection using *IRF5-* and *IKBKB-* encoding plasmids could modulate the TAM phenotype, we analyzed TAMs isolated from PDAC spheroids and pre-conditioned TAMs transfected in 2D culture with a 1:1 *IRF5:IKBKB* plasmid combination. Both transfected TAMs isolated from PDAC spheroids and pre-conditioned TAMs displayed a clear upregulation of *IRF5* accompanied by increased *IFNA* and *IFNB1*expression (Supplementary Fig. 3a-d). These findings indicate that *IRF5* and *IKBKB* transfection activates type I interferon-associated transcriptional programs in TAMs that could promote TME remodeling towards a more tumor suppressive environment in the PDAC spheroid model.

To optimize optoporation parameters in 3D, we first delivered tdTomato mRNA to TAMs in the naive AuNP^+^ system (Supplementary Fig.4). We then assessed the feasibility of TAM-transfection via optoporation in both naive AuNP^+^ and conditioned AuNP^+^ spheroids alongside with a naive AuNP^-^ spheroid as control (Fig. 3a,c,e). Spheroids were incubated with a 1:1 mixture of plasmids encoding IRF5-GFP and IKBKB-GFP and subjected to optoporation (Opto^+^) targeting CellTrace Violet (CTV)-labelled TAMs.

**Figure 3.**
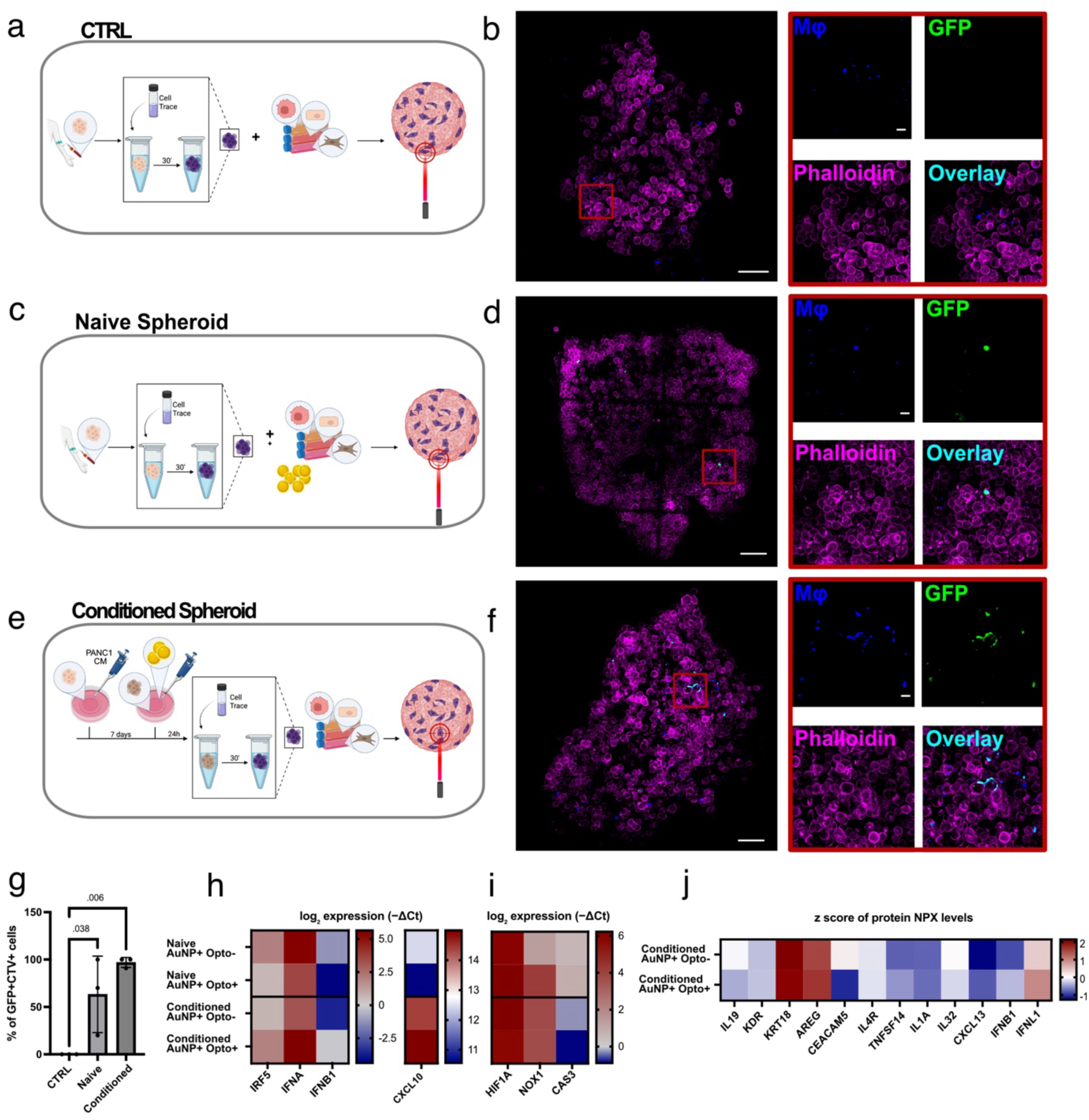
Optoporation-mediated TAM-specific delivery of *IRF5-* and *IKBKB-* plasmids drives TME remodeling in 3D PDAC spheroids. Schematics of the targeted 3D optoporation workflow in: **a,** naive AuNP^-^ spheroid (CTRL), **c,** naive AuNP^+^ spheroid and **e,** conditioned AuNP^+^ spheroid. Maximum intensity projections of confocal images of optoporated spheroids stained for F-actin (phalloidin, magenta), showing CTV-labeled macrophages (purple), GFP expression from transfected plasmids (green), and overlay. Scale bars: 100 μm.: **b,** naive AuNP^-^ spheroid, **d,** naive AuNP^+^ spheroid and **f,** conditioned AuNP^+^ spheroid. **g,** Quantification of the CTV^+^GFP^+^ cells from image analysis. **h-i,** qPCR quantification (log₂ expression, −ΔCt) of *IRF5, IFNA, IFNB1, CXCL10* (h) and *HIF1A, NOX1, CAS3* (**i**) for naïve and conditioned AuNP^+^ spheroids w/wo optoporation. **j,** Z-score of selected NPX protein levels for the conditioned AuNP^+^ spheroids w/wo optoporation. Data represent mean ± SD from three biological replicates (n=3). Statistical analysis was done by one-way ANOVA with Dunnett’s test for multiple comparisons; p ≤ 0.05 was considered significant. Schemes in **a,c,e** were created with BioRender.

At 24 h post-optoporation, both naive and conditioned AuNP^+^ spheroids showed clear colocalization of GFP expression with CTV signal, demonstrating successful plasmid transfection into targeted TAMs, while AuNP^-^ controls exhibited no GFP signal (Fig. 3b,d,f). Quantification revealed an averaged transfection efficiency of 63.33% CTV^+^ GFP^+^ cells in naive AuNP^+^ spheroids compared to an average of 96.67% CTV^+^ GFP^+^ cells in conditioned AuNP^+^ spheroids, with 0% transfected cells in monocyte-seeded AuNP^-^ controls (Fig. 3g). As expected, no signal was detected when incubating with AuNPs without laser illumination (Supplementary Fig.5).

We then evaluated whether delivery of *IRF5* and *IKBKB-*plasmids via optoporation induced functional TME modulation in the PDAC spheroids. Gene expression analysis revealed striking differences between the two spheroid models: naive AuNP^+^ Opto^+^ spheroids showed no notable changes in the selected gene expression, whereas conditioned AuNP^+^ Opto^+^ spheroids exhibited increased expression of *IRF5*, *IFNA*, *IFNB1* and *CXCL10* (Fig. 3h), analogous to TAMs transfected *in vitro* (Supplementary Fig. 3). These data indicate that targeted TAM optoporation initiates type I interferon-associated transcriptional programs within 24 h, which may contribute to a less tumor-supportive microenvironment.

To confirm that observed transcriptional changes resulted from IRF5/IKBKB signaling rather than laser-induced cellular stress, we assessed expression of hypoxia-, ROS-, and early apoptosis-associated genes. *HIF1A*, *NOX1* and *CAS3* showed minimal or absent upregulation in conditioned AuNP^+^ Opto^+^ spheroids (Fig. 3i). *NOX1* showed a modest increase in naive AuNP^+^ Opto^+^ spheroids (Fig. 3i).

To further confirm the TME remodeling observed with gene expression, we analyzed cytokines and other tumor-associated secreted factors in conditioned spheroid supernatants at 24 h post-optoporation. Comparisons were made between AuNP^+^ spheroids with and without optoporation (AuNP^+^Opto^−^ vs. AuNP^+^Opto^+^). AuNP presence alone induced a modest increase in epithelial keratin-18 (KRT18), regularly expressed by PDAC^34^ (Supplementary Fig. 6). Interestingly, targeted TAM optoporation in conditioned AuNP^+^ spheroids induced a coordinated shift in the secreted proteome consistent with IRF5-driven repolarization toward a tumor-suppressive TME. Type I interferon responses were elevated, with IFN-β1 protein levels strongly increased in conditioned AuNP^+^ Opto^+^ spheroids, in agreement with the gene expression data (Fig. 3j, Supplementary Fig. 6). Furthermore, IFN-λ1, a type III interferon associated with antitumor activity in several tumor models^35–37^, was elevated in the optoporated sample (Fig. 3j) while KRT18 levels decreased (Fig. 3j, Supplementary Fig. 6). Notably, conditioned AuNP^+^ Opto^+^ spheroids also showed a reduction in amphiregulin (AREG), associated with tumor-promoting EGFR signalling^38^ and KDR (VEGFR2), indicative of decreased angiogenic signaling^39^ (Fig. 3j). Furthermore, we observed altered concentration of cytokines commonly associated with pancreatic cancer promotion. IL-4R^40^ and IL1-A^41^ showed a subtle decrease, whereas IL-19^42^ and IL-32^43^ exhibited a more pronounced reduction (Fig. 3j). Carcinoembryonic Antigen-Related Cell Adhesion Molecule 5 (CEACAM5), a tumor marker upregulated in PDAC tissue^44^, was markedly reduced in conditioned AuNP^+^ Opto^+^ spheroids (Fig. 3j, Supplementary Fig. 6). CXCL13, associated with lymphocyte recruitment^45^, increased in conditioned AuNP^+^ Opto^+^ spheroids (Fig. 3j).

Together, these findings demonstrate that AuNP-assisted optoporation enables targeted delivery of *IRF5*- and *IKBKB*-encoding plasmids into TAMs within heterocellular PDAC spheroids, and that this TAM-specific optoporation is inducing type I interferon-driven macrophage repolarization, upregulation of lymphocyte-recruiting chemokines such as CXCL13, and down-regulation of pro-tumorigenic factors. These shifts correlate with a coordinated remodeling of the PDAC TME toward a less tumor supportive state, positioning optoporation as a functional tool to investigate TAM reprogramming and its impact on tumor-immune interactions.

## Discussion

This study establishes a tool for targeted TAM transfection within 3D PDAC spheroids, providing a versatile platform to manipulate cells in complex tumor environments. By enabling precise, TAM-specific gene delivery in a human-relevant 3D context, our work directly addresses a key bottleneck in mechanistically dissecting TAM-reprogramming strategies in a controlled setting that mimics human disease.

TAMs have been a significant target for anti-cancer therapies, with new therapeutic approaches being developed at an accelerating pace. A major focus has been on immune reprogramming of TAMs, due to their increasing relevance in cancer TME dynamics and modulating therapy resistance^4,17,46^. Approximately 200 new therapies focusing on TAM reprogramming are currently in clinical trials; however, these have yielded overall response rates of only ∼5%, partly due to an incomplete understanding of TAM biology and their functional response to the treatments^47^. In this context, our platform introduces the ability to genetically manipulate TAMs *in situ* within a multicellular PDAC spheroid, providing a means to systematically interrogate how specific reprogramming strategies alter TME state and to prioritize targets for therapeutic development. Because nanoparticle-assisted optoporation relies on physical membrane permeabilization, the same platform is, in principle, compatible with a broad range of nucleic acid cargos (i.e., plasmid DNA, mRNA or CRISPR-based reagents). A critical design consideration for such a platform is how nanoparticles are integrated within the 3D architecture, as their spatial distribution directly impacts the precision of optoporation. Although incorporating AuNPs during spheroid generation led to a predominant TAM uptake, a substantial fraction of ∼45% accumulated in the ECM, potentially inducing off target optoporation. To overcome this limitation and further refine spatial specificity, forming spheroids using AuNP-loaded TAMs comparatively yielded a more efficient optoporation and increased downstream effects (Fig. 3h,i,j), suggesting a benefit of achieving a more selective AuNP localization within TAMs in reducing background effects from other cells or matrix.

TAM-specific modulation with *IRF5-* and *IKBKB-* encoding plasmids induced robust activation of type I interferon expression (*IFNA*, *IFNB1*, *CXCL10*), accompanied by global changes in secreted factors. We observed a reduction of pro-tumorigenic IL-19, IL-32, CEACAM5, and KDR, alongside increase in IFN-β1, IFN-λ1 and CXCL13 which suggest that TAM-specific modulation impacts multiple axes of tumor support, including stemness^42^, angiogenesis^39^, and immune recruitment^45,48^. Lowered KRT18 levels could be reflecting macrophage-mediated tumor cell killing, proteolytic keratin degradation, or a shift toward keratin-low tumor cell populations. Changes in epithelial and stromal phenotypes are, in fact, likely to arise from a combination of direct effects of interferon signaling and indirect consequences of altered macrophage-tumor crosstalk that collectively, contribute to a shift toward a less tumor-supportive microenvironment. From a therapeutic standpoint, these coordinated changes imply that reprogrammed TAMs could act as microenvironmental amplifiers, converting a local genetic perturbation into system-wide remodeling of epithelial, stromal, and vascular compartments. Future studies could apply this technology to more complex *in vitro* systems incorporating functional vasculature in the TME, such as microphysiological systems (MPS), to assess how vascularization influences TAM reprogramming and paracrine signaling ^49–51^.

Our results in 3D models are consistent with previous studies showing that overexpression of *IRF5* in TAM promotes generation of an anti-tumor phenotype^15^. In ovarian and melanoma cancers, the increase in IL-1β seems to play a crucial role in the TAM response^15^. In contrast, for PDAC, the overexpression of *IRF5* in TAMs is characterized more by the type I interferon response, which aligns better with the TME of PDAC^8^. This highlights the complexity and specificity of immune responses in different cancer types, making it crucial to understand these distinctions for developing targeted therapies. Further, given that TAMs in tumor represent a highly heterogeneous population^52–54^, key future challenge will be to selectively reprogram distinct TAM subsets, determine the number and persistence of TAMs subjected to optoporation to influence the overall TME response, providing important insights into the durability and broader systemic impact of this approach.

## Conclusions

This research study demonstrates that optoporation can be effectively applied in heterocellular 3D spheroid models to enable spatially controlled manipulation of TAMs. This precise, cell-specific perturbation enables direct investigation of how changes in TAM phenotype influence the TME, positioning optoporation as a powerful tool to interrogate and reprogram immune– tumor interactions in physiologically relevant 3D tissues. By systematically delivering specific nucleic acids into TAMs, we uncover mechanistic insights into their function, cytokine signaling, and cell-cell interactions *in situ*. Such a versatile platform for testing targeted therapeutic interventions enables TAM-reprogramming strategies to be refined in human-relevant 3D systems, with the potential to de-risk macrophage-targeted clinical programs in indications where current therapeutic response rates remain low. In parallel, it deepens our understanding of TAM biology and their contributions to the TME, ultimately guiding the rational design of more effective and selective therapeutic strategies.

## Materials and methods

### Cell culture

Human pancreatic cancer cells PANC-1 (ATCC, USA; CRL-1469) were used as tumor model. PANC-1 cells were transduced to stably express green fluorescent protein (GFP). PANC-1 and GFP-PANC1 were cultured in medium containing DMEM (Gibco, Thermofisher, USA) supplemented with 10% fetal bovine serum (FBS, Gibco, Thermofisher, USA) and 1% antibiotic (penicillin/streptomycin, Gibco, Thermofisher, USA). Pancreatic stellate cells (PSC, Gene Ethics, Singapore; 3830) were kept in Stellate Cell Medium (SteCM, ScienCell Research Laboratories, USA; 5301) supplemented with 2% FBS (ScienCell Research Laboratories, USA; 0010), 1% stellate cell growth supplement (ScienCell Research Laboratories, USA; 5352), and 1% antibiotic solution (ScienCell Research Laboratories, USA; 0503). Human umbilical vein endothelial cells (HUVEC, Lonza, Switzerland; C2519AS) and RFP-HUVEC (Angio-Proteomie, USA; cAP-001RFP) were cultured in endothelial cell media supplemented with growth factors included in the EGM®-2 Endothelial Cell Growth Medium-2 BulletKit® (Lonza, Switzerland; CC-3162). All cells were cultured at 37 °C in a humidified atmosphere containing 5% CO_2_.

Peripheral blood mononuclear cells (PBMCs) were isolated from blood cones using Ficoll-Paque PLUS (GE Healthcare, USA). De-identified human blood tissue was collected from blood cones in accordance with and under the following project: HSA Residual Blood Samples for Research, project titled “Harnessing immune response for new therapies in transplantation and cancer” (A*STAR IRB Reference Number: 2024-133). Blood was diluted 1:40 in phosphate-buffered saline (PBS, Gibco, USA; 20012027) and layered over Ficoll-Paque (density 1.077 g/mL), followed by centrifugation at 900 ×g for 20 min at room temperature (RT). The PBMC layer was collected, washed with Ca²⁺/Mg²⁺-free PBS, and treated with red blood cell lysis buffer (155 mM NH₄Cl, 10 mM KHCO₃, 0.1 mM EDTA (1st Base, Singapore; BUF-1052-1L, pH 8.0)) for 5 min at RT. After washing, cells were cryopreserved in Bambanker™ (Fuji-film Wako Chemicals, USA). For experiments, thawed PBMCs were used to isolate untouched monocytes with the Pan Monocyte Isolation Kit (Miltenyi Biotec, Germany). Monocytes were further used either directly in spheroid formation, or they were pre-conditioned with PANC-1 media generating TAMs.

For TAMs differentiation, monocytes were plated in a T75 flask with 70% of filtered PANC-1 conditioned media and 30% of normal growth media containing RPMI 1640 (Gibco, Thermofisher, USA) supplemented with 10% FBS and 1% penicillin/streptomycin, and kept at 37 °C in a humidified atmosphere containing 5% CO_2_ for 7 days with one media change on day 4.

### Spheroid formation

Spheroids were generated using a hanging-drop method with a custom PDMS mold aligned over 96-well plates. Each spheroid contained 1500 PANC-1 cells, 3000 PSCs, 3000 HUVECs, and 6000 monocytes or TAMs. In specific experiments, fluorescently labeled cells were used to visualize individual populations: GFP-expressing PANC-1 cells, CMRA-labeled PSCs (Thermo Fisher Scientific, USA; C2927), RFP-expressing HUVECs, and CellTrace™ Violet– labeled monocytes (CTV; Thermo Fisher Scientific, USA; C34557). Spheroids were prepared under four conditions: (i) without gold nanoparticles (AuNPs), (ii) with 6 µg/mL AuNPs (Nanopartz, USA; A11-200-CIT-DIH-1-25-CS) or fluorophore-tagged AuNPs (Nanopartz, USA; CF11-200-AF680-FPEG-DIH-50-1), (iii) incorporating TAMs, or (iv) incorporating TAMs pre-incubated with 6 µg/mL AuNPs for 24 h (TAM-AuNP). Each spheroid was formed in a 35 µL drop of EGM®-2 Endothelial Cell Growth Medium-2 BulletKit® (Lonza, Switzerland; CC-3162) and maintained at 37 °C in a humidified atmosphere containing 5% CO₂ for 7 days.

### Cell viability assay

For the cell viability assay, cells were seeded in 96-well plates (Thermofisher, USA) at 40x10^3^ cells/cm^2^ (or for spheroids, one spheroid was seeded per well). Viability was determined using ATP assay CellTiter-Glo 3.0 (Promega, UK; G9241) according to the manufacturer’s instructions. Cells were incubated with gold nanoparticles for 3 h. In the case of spheroids, spheroids were formed in the presence of gold nanoparticles. Following incubation, an equal volume of reagent was added to the volume of cell culture medium present in each well (1:1 ratio) followed by incubation at 37 °C for 1 h. Solution was transferred to a dark 96-well plate and absorbance was measured using BioTek Synergy H1 Multimode microplate reader (Agilent, USA) with endpoint luminescence and gain of 200.

### mRNA extraction and qRT-PCR

At least 3 spheroids (or three wells for 2D experiments) from 3 different experiments were pooled together for gene expression analysis. Total RNA was harvested using the PicoPure RNA isolation kit (Thermofisher, USA; KIT0204), according to the manufacturer’s instructions. NanoPhotometer NP80 (Implen, Germany) was used to measure the concentration of the extracted RNA and to assess RNA quality and purity. Measurements were subjected to quality control minimum standards of A260/230 > 2 for quality and A260/280 > 1.8 for purity. First strand complementary DNA (cDNA) was synthesized from 3 μg of the total RNA, using the iScript cDNA Synthesis Kit (Bio-Rad, UK; 170-8891) according to manufacturer’s protocol. qRT-PCR was performed using a CFX384TM Real-Time System C1000TM Touch Thermal Cycler (Bio-Rad, UK) and POWER SYBR Green master mix (Thermofisher, USA). Gene expression levels were quantified using the comparative threshold cycle (CT) method, normalized to the reference gene HPRT1 and expressed as 1/(2^ΔCT^), where ΔCT is the difference between the CT of the target gene and that of the reference genes. Primer sequences are listed in Table 1.

**Table 1.**
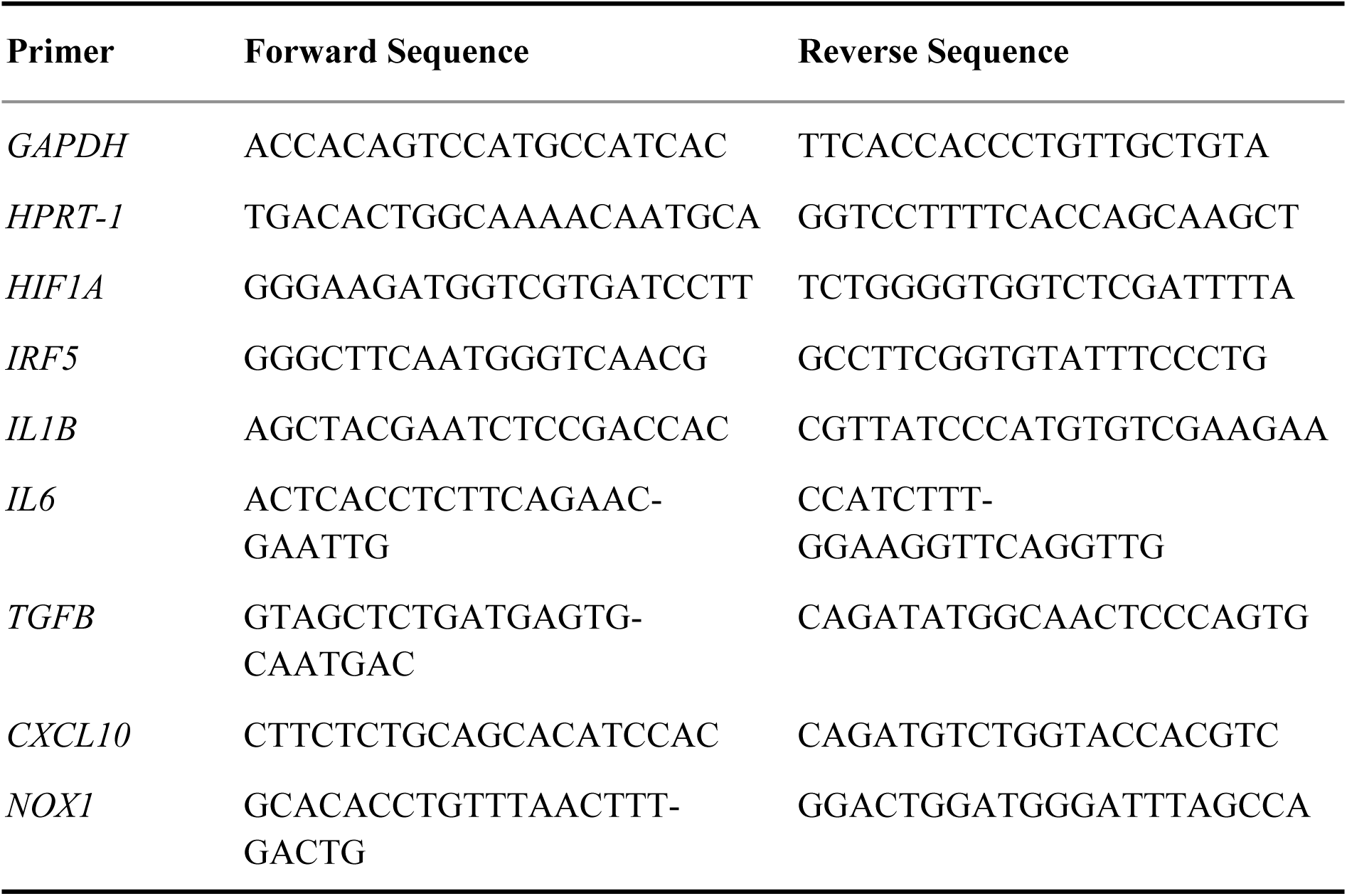
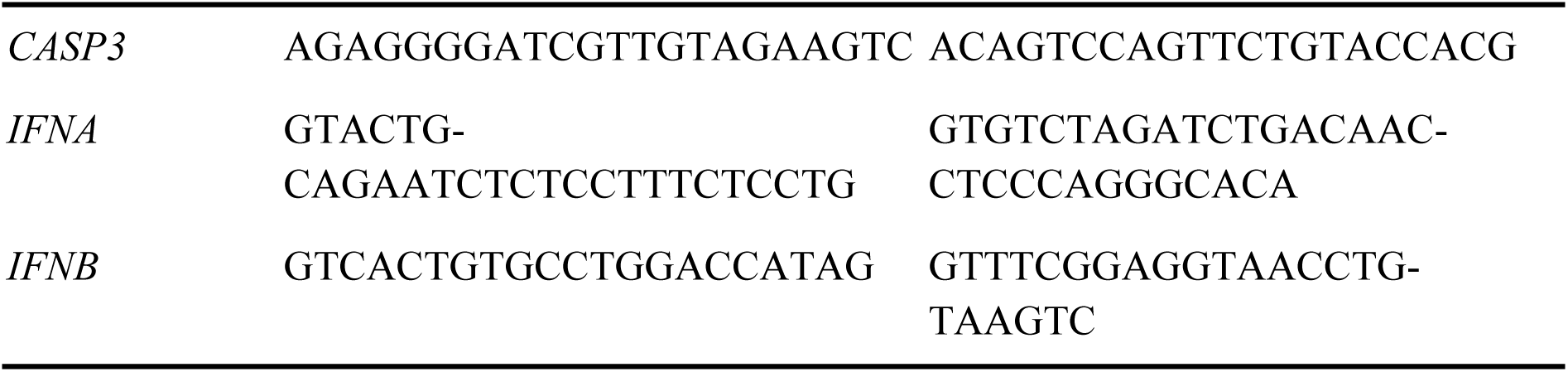
Primer sequences.

### Immunofluorescence microscopy

3D Spheroids: Spheroids were collected and fixed with 200 µL of 4% paraformaldehyde (Thermo Scientific, USA; J19943-K2) at RT for 1 h. Afterwards, they were washed twice with PBS. Spheroids that were stained with immunofluorescent antibodies were then permeabilized with 0.25% Triton X-100 in PBS for 15 min and blocked with 3% BSA (Bio-Rad, UK; 5000206) and 2% Donkey Serum (Sigma Aldrich, Germany; D9663) in PBS for 1 hat RT. After washing, the spheroids were incubated overnight at 4 °C with primary antibody against Pan Collagen (1:200, ThermoFisher, USA; PAl-55324). After washing, the spheroids were then incubated with secondary antibody (1:1000 donkey anti-rabbit DyLight488, BioLegend, USA; 406404) and Hoechst 33342 for nuclei visualization (1:1000; ThermoFisher, USA; HI399) for 6 hat RT or phalloidin AF647 for cellular actin filaments (1:200, ThermoFisher, USA; A22287) for 15-30 min. Pre-labeled cells (GFP-PANC-1, RFP-HUVECs, CMRA-PSC, CTV-monocytes/TAMs) were used where indicated.

2D cultures: Post-differentiated TAMs were incubated with 6 µg/mL fluorophore tagged AuNPs (Nanopartz, USA; CF11-200-AF680-FPEG-DIH-50-1) and labeled with CellTrace™ Green (Thermo Fisher Scientific, USA; C2925), nuclei counterstained with Hoechst 33342 (1:1000, ThermoFisher, USA, H1399). Lysosomes were labeled using CellLight Lysosomes-GFP BacMam 2.0 (Invitrogen, USA; C10596) according to the manufacturer’s instructions, followed by incubation under standard culture conditions prior to imaging. Fixation, permeabilization, and blocking were performed as for spheroids.

Images were acquired using Olympus IX83 Inverted Microscope System (Olympus, Japan) with a 20X (NA 0.8, air) objective using the Olympus FV31S-SW Software (Olympus, Japan) and analyzed with Fiji ImageJ software (Version 2.14.0, National Institutes of Health, USA).

### Flow cytometry

3D spheroids: At least 10 spheroids were pooled in a microtube, washed with PBS, and dissociated in 50 µL of 5 mg/mL Collagenase IV (Gibco, USA; 17104019) and TrypLE™ 1X Express (Gibco, USA; 12604013) for 20–30 min at 37 °C, with pipetting every 5 min to facilitate dissociation into single cells. Cells were washed twice with PBS (1500 rpm, 5 min, RT) and resuspended in 100 µL of staining buffer (MACS buffer: 1X PBS, 0.5% BSA, 2 mM EDTA containing the relevant antibodies. Samples were incubated for 30 min at RT on a shaker, then washed twice with PBS and resuspended in staining buffer for acquisition.

2D cultures: Monocytes and post-differentiated TAMs were incubated with 6 µg/mL fluorophore-tagged AuNPs (Nanopartz, USA; CF11-200-AF680-FPEG-DIH-50-1) for 0 and 3 h (monocytes) or 0, 3, and 24 h (TAMs). After incubation, cells were washed once with PBS, stained with Calcein-AM to assess viability, washed again, and treated with TrypLE™ 1X Express (Gibco, USA; 12604013) for 10 min prior to acquisition.

All experiments were analyzed by flow cytometry (BD Symphony). Experiments were performed at least in triplicate, and 10000 cells per sample were measured. Antibodies used are listed in Supplementary Table 2.

### Lipofectamine transfection

Post-dissociation, CD14⁺ cells were isolated from standard (non–TAM-bearing) spheroids using magnetic beads (Miltenyi Biotec, Germany; 130-097-052). Preconditioned TAMs were collected and plated directly in 3 wells per experiment. For CD14⁺ cells, 180 spheroids were generated per experiment, cells were extracted and transfected, and this procedure was repeated in 3 independent experiments. TAMs were transfected following the same procedure for 3 independent experiments. Cells from all replicates were lysed in PicoPure Extraction Buffer (Thermo Fisher Scientific, USA; KIT0204) and pooled per condition for downstream gene expression analysis.

For transfection, cells were incubated with Lipofectamine 3000 (Thermo Fisher Scientific, USA; L3000001), prepared according to the manufacturer’s instructions, and plasmids IRF5 (OriGene, USA; RG200458) and IKBKB (OriGene, USA; RC219154L4) at a 1:1 ratio (total concentration 0.3 µg/mL) for 48 h under standard culture conditions. Following incubation, cells were harvested for qRT-PCR analysis.

### Laser treatment

All experiments (2D TAM-AuNP and 3D spheroids) were performed using the Nikon Eclipse Ti with Apo WI l S DIC N2, N.A 1.25 (Nikon, Japan) and Spectra-Physics Mai Tai laser system (Spectra-Physics, USA), delivering <100 fs near-infrared pulses at 800 nm with an 80 MHz repetition rate. Samples were irradiated from below with a focused beam on the targeted area. Laser power was set as a percentage of the total output power displayed in the NIS-Elements software (≈2900 mW after stabilization) for the Spectra-Physics Mai Tai. The applied powers therefore correspond to estimated average output powers of 145 mW (5%), 203 mW (7%), 222 mW (7.5%), and 442 mW (15%).

3D spheroids: Spheroids (naive or conditioned AuNP^+^) containing CellTrace™ Violet–labeled monocytes/TAMs (CTV; Thermo Fisher Scientific, USA; C34557) were plated into 50 µm glass-bottom dishes (Ibidi, USA; 81148), one spheroid per well in a culture insert (Ibidi, USA; 80366). Following plating, they were washed one time and media was changed to RPMI 1640 phenol free (Gibco, Thermofisher, USA) supplemented with 2% heat inactivated FBS. Immediately before laser irradiation, a 1:1 mixture of IRF5 (OriGene, USA; RG200458) and IKBKB (OriGene, USA; RC219154L4) plasmids, prepared at a total concentration of 0.3 µg/mL, was added. Each treatment lasted 4 s per frame per area delivering 222W of power. Negative controls (laser irradiation without AuNPs) were included.

2D TAM-AuNP optimization: Post-differentiated TAMs incubated with AuNPs for 24 h were plated on glass-bottom dishes, and prior to irradiation, media was replaced with RPMI 1640 phenol-free (Gibco, Thermo Fisher Scientific, USA) supplemented with 2% heat-inactivated FBS. Propidium iodide (Miltenyi Biotec, Germany; 130-093-233) was added before irradiation to assess cell membrane integrity, and Calcein-AM (Thermofisher, USA, C1430) was applied post-irradiation to evaluate viability. Cells were irradiated under multiple conditions to optimize parameters for plasmid delivery, with power of 145W, 203W, 222W and 442W and scanning patterns of 4 and 6 spatial frames per area (SPFs). Negative controls (no AuNPs) were included in all conditions.

Post-irradiation, spheroids and/or 2D cultures were imaged using an Olympus IX83 inverted microscope (20x air objective, NA 0.8 or 40x air objective, NA 0.95) with Olympus FV31S-SW software (Olympus, Japan) and analyzed using Fiji/ImageJ (v2.14.0, NIH, USA). Additionally, at least one spheroid per experiment was collected in PicoPure Extraction Buffer (Thermofisher, USA; KIT0204) for subsequent qRT-PCR analysis.

### Single cell analysis

scRNA-seq was performed on PDAC spheroids as described in^29^. Briefly, 60 PDAC spheroids (per condition) were collected 7 days after seeding and dissociated using collagenase IV followed by trypsin/EDTA digestion at 37 °C. Enzymatic activity was stopped with EGM2, and cells were washed with PBS + 1% BSA. Single-cell suspensions were processed on the 10x Genomics Chromium Controller (target: ∼6000 cells/sample) for library preparation using the Chromium Next GEM Single Cell 3′ v3.1 Kit. Libraries were sequenced on an Illumina NovaSeq 6000 (2×151 cycles, ∼50000 reads/cell). This work analyzes only one of the four culture conditions sequenced, relative to the monocyte-seeded PDAC spheroids.

Raw count matrices were imported into R (v3.18) and analyzed with Seurat (v5.0.3)^55,56^. Low-quality and doublet cells were removed based on number of genes detected (more than 200 and less than 7000) and mitochondrial gene expression (with less than 20 % mitochondrial gene expression). Data was then normalized using LogNormalize method. Variable genes were identified with variance-stabilizing transformation and top 2000 genes were chosen for the further downstream analysis. Following with clusters identification and dimensionality reduction conducted using UMAP with 17 dimensions based on elbow plot. To further resolve heterogeneity within the macro-phage-like cluster, we performed supervised subclustering. A subset of macrophage cells was extracted from the main Seurat object, and the expression of selected marker genes (*CCL2, IL1B, APOE, CD163, CD40, CD274, STMN1, CD14, CD52*) was scaled and used to perform principal component analysis. The top three principal components were used to construct a shared nearest neighbor graph, followed by clustering with a resolution of 0.5 and UMAP di-mensionality reduction for visualization. The resulting subcluster assignments were added to the main Seurat object for downstream analyses. This approach enabled identification of functional-ly distinct macrophage subpopulations within the 3D spheroid model. Differential gene expression (DEG) was per-formed using FindAllMarkers function with a fold change threshold of 0.25, an adjusted p-value threshold of 0.1 and false discovery rate (FSR) < 0.05 considered as statistical significance. Furthermore, we performed enrichment analysis using ClusterProfiler (v4.10.1) ^57,58^. EnrichGO function including molecular function (MF), biological process (BP) and cellular component (CC). Visual representations of the data were generated using Seurat or ClusterProfiler internal functions or ggplot for heatmaps.

### Protein analysis

Protein expression was measured using the Olink® Explore platform (Uppsala, Sweden) with the Olink Target 48 Immune Surveillance panel, following the manufacturer’s protocol. Briefly, 3 samples were pooled together and incubated with antibody-oligo conjugates, followed by PCR amplification and Illumina sequencing. Raw digital counts were converted to NPX (Normalized Protein eXpression) values. Data was normalized across experimental batches using the Olink® Analyze R package (version 4.3.0). Subsequent statistical analysis, including differential expression testing (e.g., Welch’s t-test) was performed in R (v3.18) to identify significantly altered proteins (adjusted p-value < 0.05). Data visualization, including heatmaps and volcano plots, was generated using the same package.

### Statistical analysis

Statistical analyses were performed using GraphPad Prism 9. Data are represented as mean ± standard error of mean (SEM) from 3 replicates and of at least 3 independent experiments, unless otherwise stated. Normal distribution of the data was tested using the Shapiro-Wilk test. One-way ANOVA or student t-test was used to test differences between groups on the normally distributed data. Testing for statistical differences between variables that did not follow normal distribution were determined with Kruskal-Wallis H test or Mann-Whitney test. p≤0.05 was considered statistically significant.

## Acknowledgments

Image acquisition was performed at the A*STAR Microscopy Platform (AMP). The authors thank the AMP and the AMP staff at the Skin Research Institute of Singapore, especially Xiaoxiao Ma, PhD and Shuping Lin, PhD, for technical support and assistance with confocal imaging. Flow cytometry was performed at the A*STAR Singapore Immunology Network (SIgN) Flow Cytometry Platform. The authors thank the staff at the SIgN Flow Cytometry Platform, especially Ivy Low and You Yi Leon Hwang, PhD, for technical support and assistance with flow cytometry experiments and data acquisition. Olink analysis was performed at Viral Research and Experimental Medicine Centre at SingHealth Duke-NUS (ViREMiCS). The authors thank the ViREMiCS staff, especially Valerie Shyn Yun Chew and Ong Ziying Eugenia, PhD, for technical support with Olink assay.

## Ethical approvals

De-identified human blood tissue was collected from blood cones in accordance with and under the following project: HSA Residual Blood Samples for Research, project titled “Harnessing immune response for new therapies in transplantation and cancer” (A*STAR IRB Ref. No. 2024-133).

## Contributions

IP performed experiments, data acquisition and analysis, contributed to methodology and project administration, wrote the original manuscript; INBH contributed to experimental work; CG contributed to experimental work; GG conceived the naive quadri-culture spheroid model, provided samples for scRNA-seq, and secured the funding that supported part of the work in SIgN; TX assisted with operating the laser; YT contributed methodology and resources; CC and GA conceived the study, secured funding, provided resources, developed methodology, oversaw project administration and supervision, and revised and edited the manuscript. All authors reviewed and approved the final version of the manuscript.

