## Supplementary material for "Immune reprogramming of 3D tumor models via optoporation-mediated targeted gene delivery to macrophages": Suplemental Material

This PDF file includes:

Supplementary Tables 1 and 2.

Supplementary Figures 1 to 6.

Other supplementary materials (provided as separate files):

Supplementary Data File S1 to S4.

| Delivery parameter | Optimal Condition |
|---|---|
| Pulse energy | 2.78 nJ |
| Repetition rate | 80 MHz |
| Scan speed | 4 seconds/frame |
| Diameter lens | 9.41 mm |
| ROI size | W 38.5 x H 28.9 µm |

**Supplementary Table 1. Overview of the optoporation conditions applied post-optimization.**

| Antibody target | Company | Fluorophore | Cat # |
|---|---|---|---|
| EpCAM | BioLegend | AF488 | 324209 |
| CD14 | BioLegend | Pacific Blue | 325616 |
| CD45 | BD | BUV395 | 563792 |
| CD31 | BioLegend | BUV605 | 303121 |
| PDGFRA | BioLegend | PE | 323505 |

**Supplementary Table 2. List of antibodies used for flow cytometry analysis.**

a

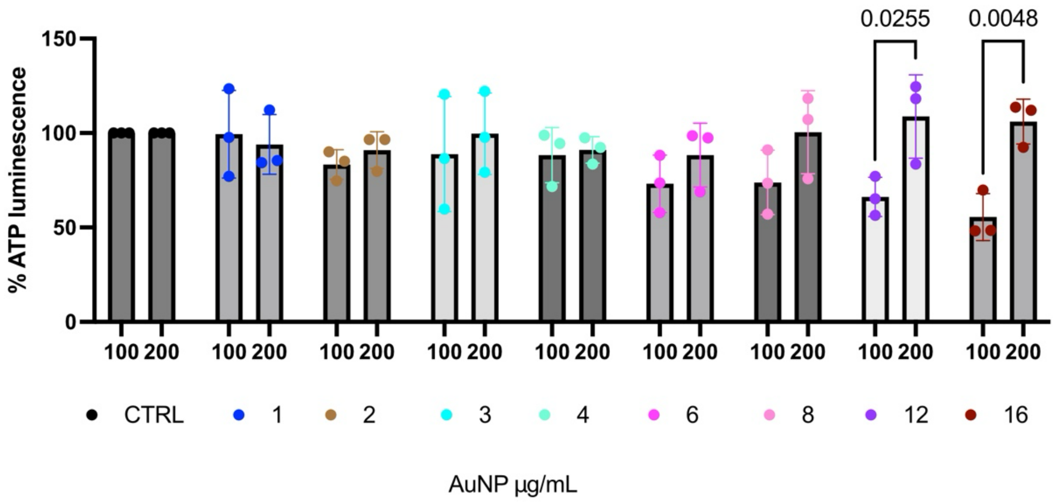


b

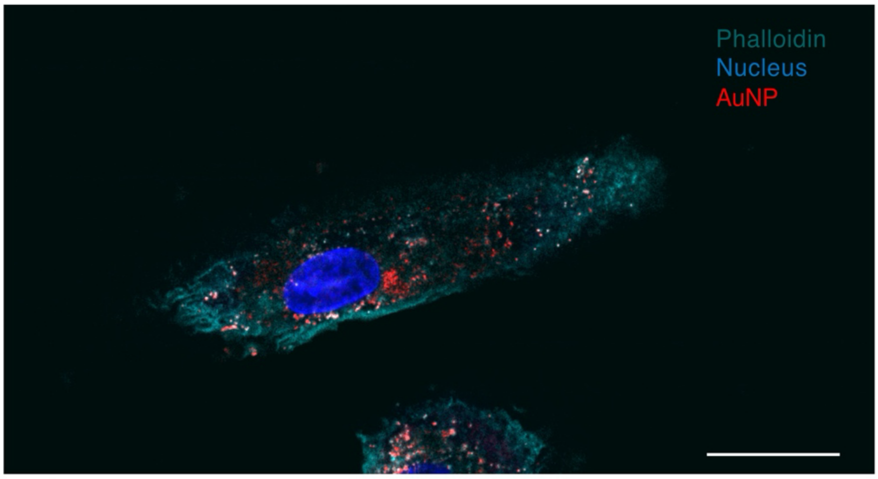


**Supplementary Figure 1. a**, Quantification of ATP-based cell viability in TAMs following 3 h incubation with or without (CTRL) AuNPs, plotted as a comparison between 100 and 200 nm AuNPs at increasing concentration. Data represent mean ± SD from three biological replicates (n=3). Statistical analysis was done by one-way ANOVA with Dunnett's test for multiple comparisons; $p \leq 0.05$ was considered significant. **b,** Representative Airyscan image of a TAM with 6 µg/mL AuNPs. Scale bar: 5 µm.

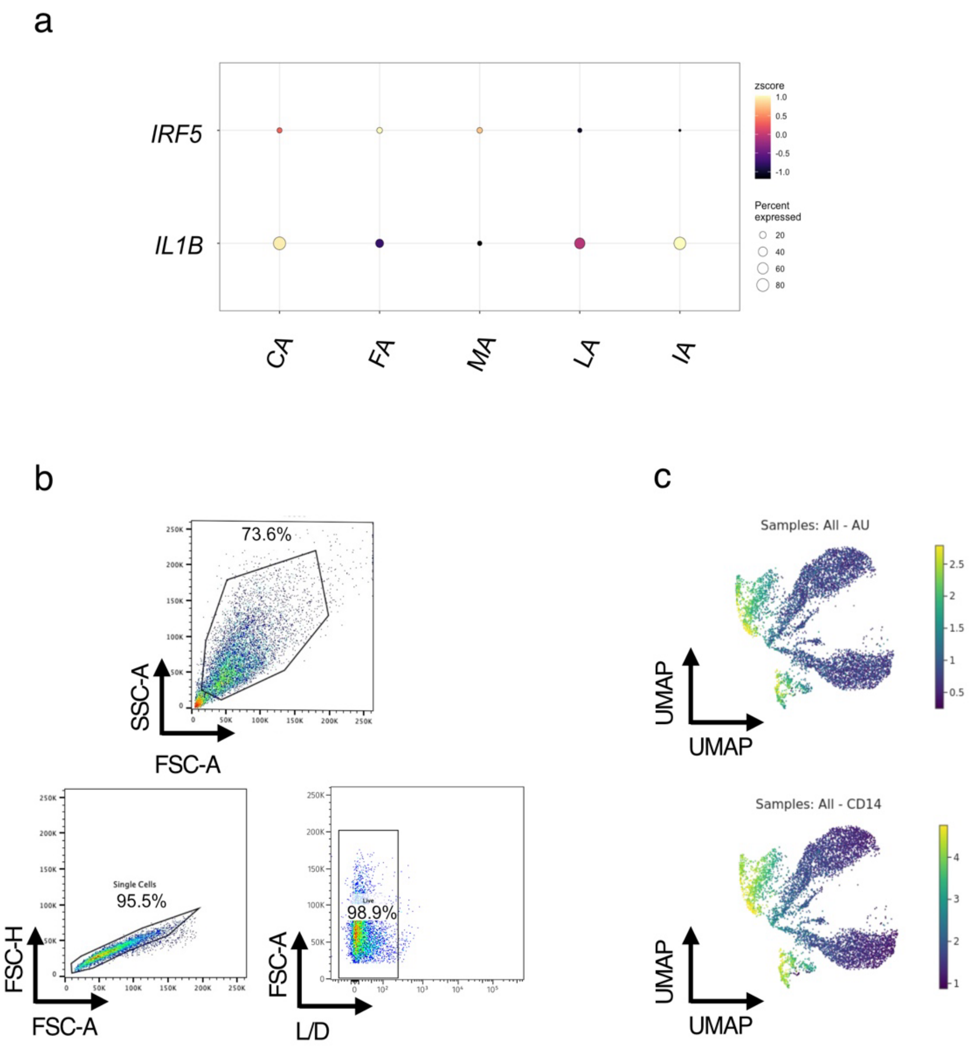


**Supplementary Figure 2. a**, Dot plot of *IRF5* and *IL1B* expression in macrophage subclusters from single cell data of classical spheroid. **b**, Gating strategy for flow cytometry data in Fig. 2.k. **c**, UMAPs of AuNP+ and CD14+ cells of flow cytometry data in Fig. 2k, Cytolytics software (Cytolytics, Germany).

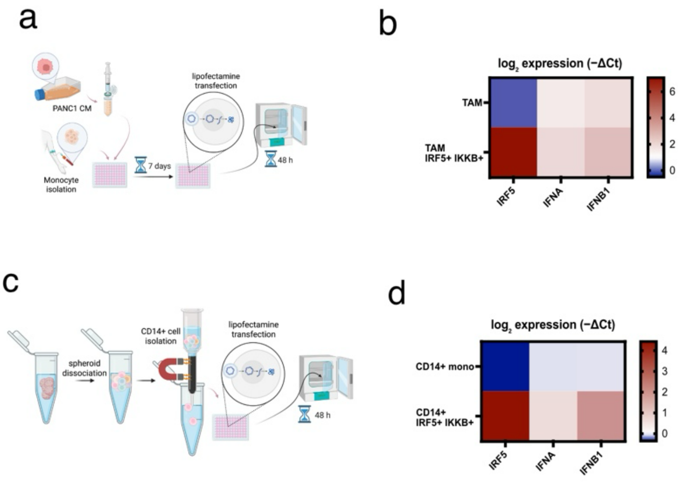


**Supplementary Figure 3**. 2D transfection of TAMs with *IRF5* and *IKBKB* plasmids. **a,** Scheme for *in vitro* TAM transfection. **b**, $\log_2$ expression (−ΔCt) of *IRF5, IFNA, IFNB1* for TAMs w/wo *IRF5/IKBKB* transfection. **c,** Scheme for *in vitro* transfection of CD14+ isolated cells from spheroids. **d**, $\log_2$ expression (−ΔCt) of *IRF5, IFNA,* IFNB1 for CD14+ isolated cells from spheroids w/wo *IRF5/IKBKB* transfection.

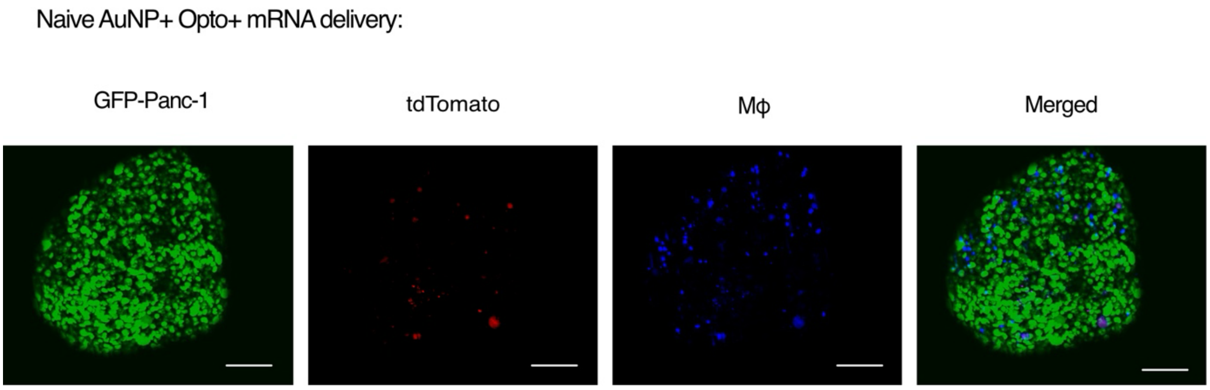


**Supplementary Figure 4.** Optoporation-mediated TAM-specific delivery of tdTomato mRNA in 3D PDAC spheroids. Maximum intensity projections of confocal images of optoporated spheroids, showing GFP-Panc-1 cancer cells (green), CTV-labeled macrophages (purple), tdTomato expression from transfected mRNA (red), and overlay. Scale bars: 100 µm.

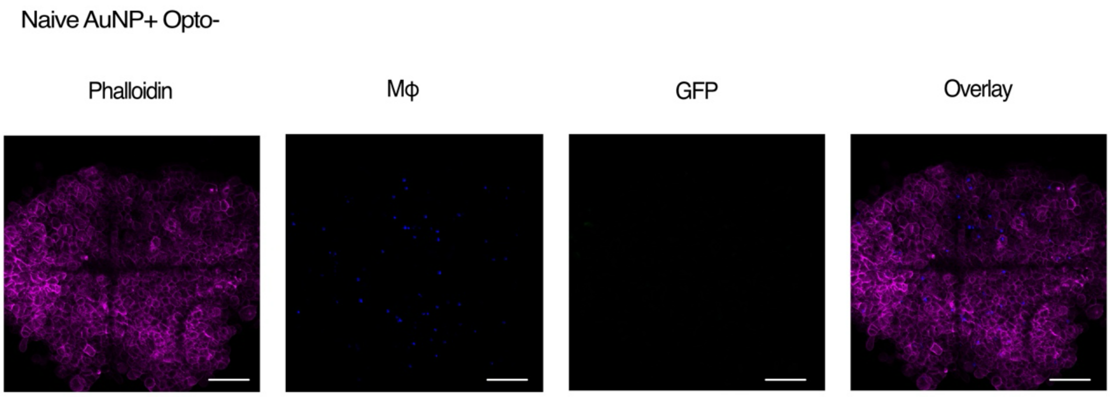


**Supplementary Figure 5.** Control showing naive spheroid incubated with AuNPs and plasmid in the absence of laser illumination, no transfection occurs. Maximum intensity projections of confocal images of optoporated spheroids stained for F-actin (phalloidin, magenta), showing CTV-labeled macrophages (purple), GFP expression from transfected plasmids (green), and overlay. Scale bars: 100 µm.

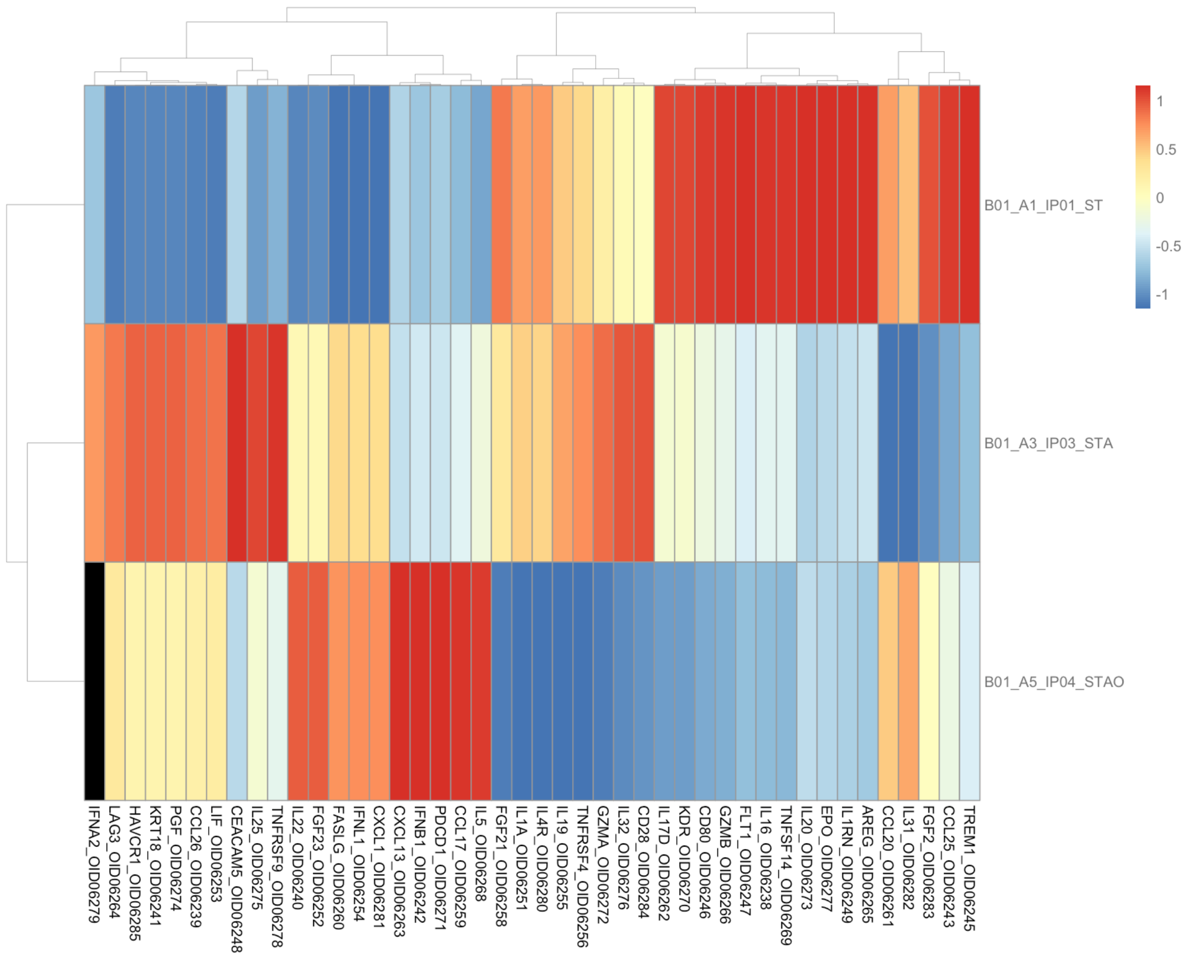


**Supplementary Figure 6.** Heatmap NPX protein levels for the conditioned AuNP+ spheroids w/wo AuNPs and w/wo optoporation. Heatmap colors are scaled per protein across all samples; NPX values are centered and scaled to facilitate visualization, with hierarchical clustering applied to both proteins and samples. Heatmap generated using Olink® Analyze Vignette. Legend: ST = AuNP- Opto-, STA = AuNP+ Opto-, STAO = AuNP+ Opto+.

**Supplementary Data File S1 (separate file)**

Gene Ontology enrichment results for biological process (BP) for the gene sets analyzed in this study.

**Supplementary Data File S2 (separate file)**

Gene Ontology enrichment results for biological process (BP), cellular component (CC), and molecular function (MF) for the gene sets analyzed in this study.

**Supplementary Data File S3 (separate file)**

Normalized protein expression (NPX) values from the Olink proteomics panel for all samples.

**Supplementary Data File S4 (separate file)**

Top differentially expressed genes for each macrophage subcluster identified in the single-cell analysis.
